# Phylogenetic Diversity and Venom Gland Transcriptomics of Palauan Cone Snails (*Conidae*)

**DOI:** 10.1101/2025.10.07.680929

**Authors:** Hudson Judge, Ho Yan Yeung, Bailey W. Miller, Zhenjian Lin, Louis R. Barrows, Patrick L. Colin, Eric W. Schmidt, Helena Safavi-Hemami, Thomas Lund Koch

## Abstract

Cone snails (genus *Conus* Linnaeus, 1758) represent one of the most species-rich lineages of venomous marine snails, producing diverse and highly specialized toxins. While *Conus* diversity is particularly high in the Indo-Pacific, some regions, such as the archipelago of Palau, remain poorly studied. Here, we combine mitochondrial marker gene sequencing (COI, 12S, and 16S) and venom-gland transcriptomics to investigate the phylogenetic relationships and venom gene diversity of Palauan *Conus* species. We sampled 27 species, recovering mitochondrial sequences for 34–41 individuals per gene. Maximum-likelihood phylogenetic analysis revealed that most individuals group with known species, although several lineages show notable divergence from reference sequences. We additionally sequenced venom gland transcriptomes from 21 individuals representing 14 species, including three species for which no previous venom data exist. Comparative analysis showed largely consistent expression patterns at the superfamily level within species collected from different biogeographical regions, supporting conserved venom gene expression profiles. These results contribute foundational data for future studies of *Conus* evolution, biogeography, and toxin diversification in an underexplored biodiversity hotspot.

## Introduction

The marine gastropod family Conidae, in particular the clade *Conus* Linnaeus, 1758, represents one of the most species-rich and ecologically diverse lineages of predatory neogastropods. Today, the *Conus* comprises 834 recognized species (World Register of Marine Species). Cone snails inhabit tropical and temperate seas worldwide, from intertidal zones to depths exceeding 1,000 meters, and are renowned for their sophisticated venom and the remarkable diversity of peptide toxins they produce. Cone snails inhabit tropical and temperate seas worldwide, with most species occurring in shallow waters from intertidal zones to the continental shelf, and some extending into deep waters at depths exceeding 1,000 meters. Each *Conus* species synthesizes hundreds of conotoxins, small, disulfide-rich peptides that target ion channels, receptors, and other proteins in their prey (Olivera *et al*., 2014). Conotoxins can be classified into approximately 70 gene superfamilies based on similarity of their signal sequences, and large-scale transcriptomic analyses have broadly validated these groupings (Robinson & Norton, 2014). While overall venom composition correlates strongly with diet (mollusks, worm, or fish-feeding), it exhibits dramatic divergence even among closely related species (Koch *et al*., 2024). Despite this diversity, many species and lineages remain poorly characterized at both the genetic and transcriptomic levels. The Indo-Pacific is a recognized biodiversity hotspot for cone snails, but the diversity of cone snail species in the archipelago of Palau (586 islands in Micronesia) remains remarkably understudied. Palau’s coral reefs and lagoon systems host a range of ecological niches that may promote cryptic speciation, local endemism, and adaptive divergence in venom composition. However, few studies have surveyed cone snail diversity in this region.

In this study, we combine mitochondrial marker gene sequencing (COI, 12S, 16S) and venom-gland RNAseq to resolve the phylogenetic relationships among *Conus* species collected from different regions in Palau and to compare their venom composition to conspecifics from other parts of the Indo-Pacific.

## Materials and Methods

### Field collections

We collected *Conus* specimens by snorkeling and scuba diving in Palau in December 2023 (Republic of Palau Marine Research Permit #RE-23-13; State Permits from Aimeliik, Airai, and Ngeremlengui). The collection localities of all specimens used in molecular studies are given in **Figure 1** and **Table 1**. Specimens were photographed, and tissues samples were preserved in RNAlater or snap frozen. Foot tissues were used for mitochondrial marker gene sequencing, while venom glands were used for RNAseq experiments.

**Table 1.** List of collected species at the different sites.

|  | *A* | *B* | *C* | *D* | *E* | *F* | *G* |
| --- | --- | --- | --- | --- | --- | --- | --- |
| *C. capitaneus* |  |  |  |  |  | X | X |
| *C. distans* | X |  | X |  |  | X |  |
| *C. ebraeus* | X |  | X |  |  |  | X |
| *C. eburneus* |  |  |  |  |  | X |  |
| *C. ferrugineus* |  |  |  |  |  | X |  |
| *C. flavidus* |  |  | X |  |  | X |  |
| *C. frigidus* |  |  |  | X |  |  |  |
| *C. imperialis* |  | X | X |  |  |  |  |
| *C. leopardus* |  |  |  | X |  | X |  |
| *C. litterarus* |  |  |  | X |  |  |  |
| *C. magus* | X |  |  |  | X |  | X |
| *C. marmoreus* | X | X |  |  | X | X |  |
| *C. miles* |  |  | X |  |  |  |  |
| *C. miliaris* | X |  | X |  |  | X | X |
| *C. moreleti* |  |  | X |  |  |  |  |
| *C. muriculatus* |  |  |  |  |  |  | X |
| *C. musicus* | X |  | X |  |  |  |  |
| *C. mustelinus* |  |  |  |  | X |  | X |
| *C. nux* |  |  | X |  |  |  |  |
| *C. pulicarius* | X |  |  |  | X | X | X |
| *C. rattus* | X |  | X |  |  |  |  |
| *C. sanguinolentus* |  |  |  |  |  | X |  |
| *C. sponsalis* |  |  | X | X |  |  | X |
| *C. striatus* |  |  |  |  |  | X |  |
| *C. textile* |  |  |  |  |  | X |  |
| *C. virgo* |  |  |  |  |  | X |  |

**Figure 1.**
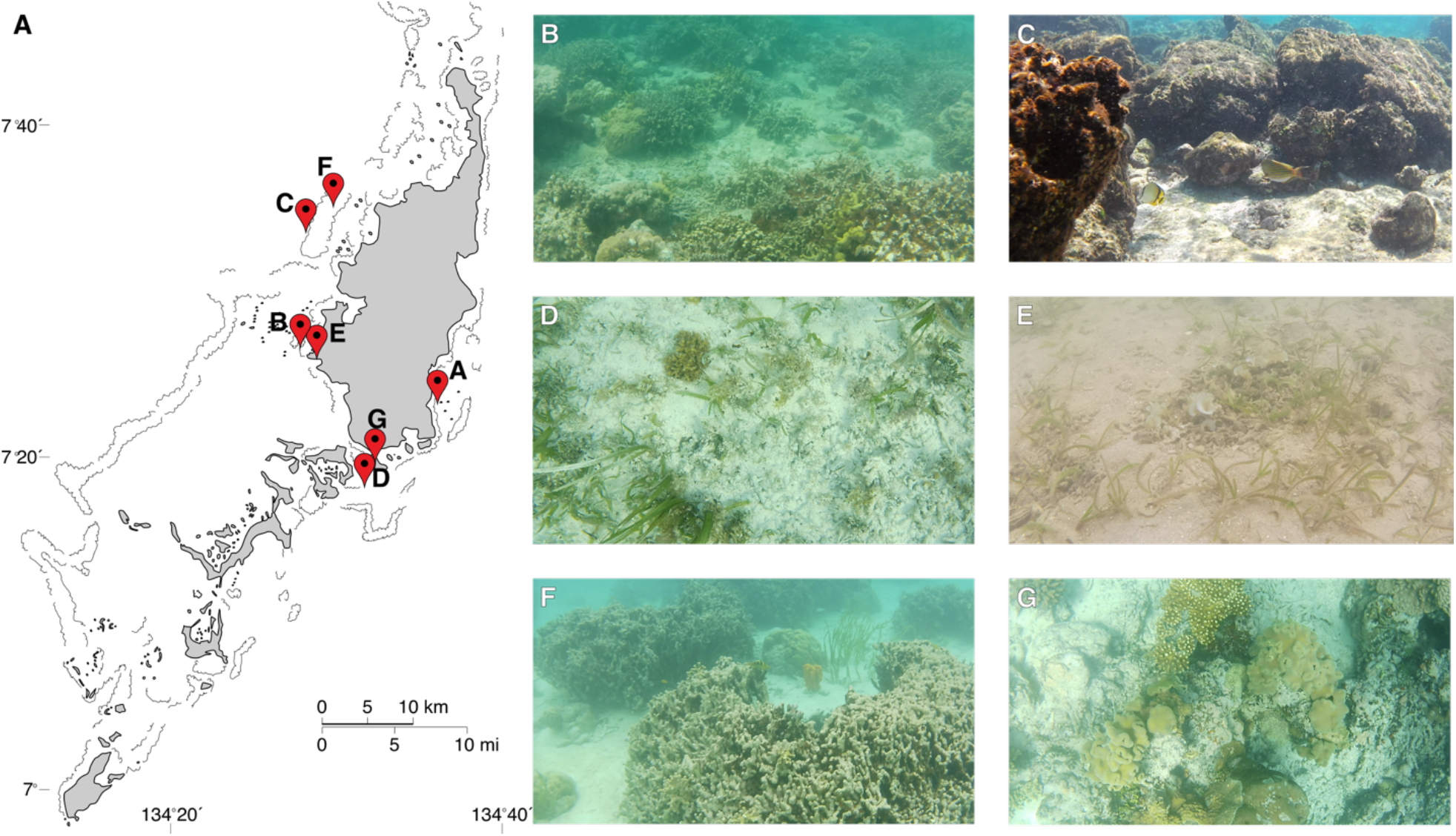
(**A**) Map showing the collection sites. Habitat photos from collection sites B-G. Site A were used for collection at night.

**Figure 2.**
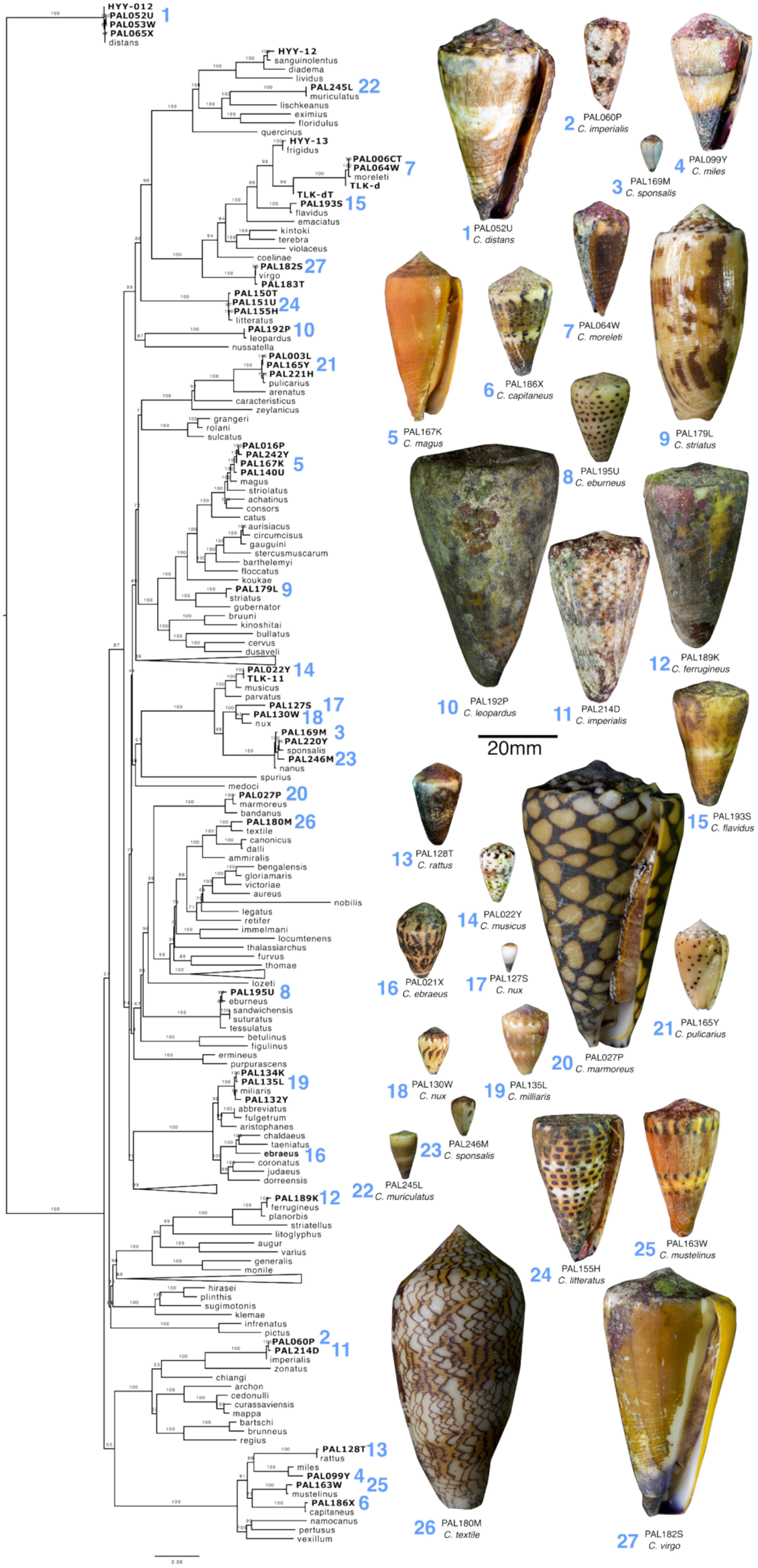
Species tree of collected *Conus* species. Maximum likelihood reconstruction of species tree of collected *Conus* species using published sequences as references. The tree was outgrouped with *C. distans*. The model of evolution was calculated with ModelFinder to be TVM+F+I+R6 for the 12S and 16S partitions and GTR+F+I+G4 for COI. The numbers show bootstrap values calculated with 1000 replicates using UF-boot.

### Mitochondrial marker gene sequencing

Approximately 10 mg of foot tissue was used for genomic DNA extraction using E.Z.N.A Mollusc and Insect DNA Kit (Omega biotek) according to the manufacturer’s protocol. Ten nanograms of DNA was used as template for polymerase chain reaction (PCR) with three primer sets targeting 12S (Forward: 5′ TCG CAG CAG YCG CGG TTA, Revers: 5′ AGA GYG RCG GGC GAT GTG T) adapted from(Hewitt, Johnston & Young, 2013), and 16S (Forward: 5′ CCG GTC TGA ACT CAG ATC ACT G, Reverse: 5′ GTT TAC CAA AAA CAT GGC TTC)(Palumbi, 1996), and mitochondrial COI (Forward: 5′ GGT CAA CAA ATC ATA AAG AYA TGY G, Reverse: 5′ TAA ACT TCA GGG TGA CCA AAR AAY CA 3′). The PCR cycling profile was as follows: Initial denaturation (94°C, 5 minutes); followed by 40 cycles of denaturation (95°C, 20s); annealing (55°C, 20s) and extension (72°C, 30s). The PCR products were run on a 1.5 % agarose TAE gel, and the desired band for the PCR product was cut and cleaned with QIAquick PCR and Gel Cleanup Kit (Qiagen) according to the manufacturer’s protocol. The cleaned-up product was sequenced at the DNA Sequencing Core at the University of Utah on an Applied Biosystems 3730xl sequencer. Nucleotide sequences sequenced in this study were submitted to NCBI (accession numbers pending).

### Venom gland transcriptomics

RNA was extracted from the venom glands using Direct-zol RNA extraction kit (Zyme Research, Irvine, CA, USA) following the manufacturer’s protocol. Extracted RNA went through library preparation with the Illumina TruSeq Stranded mRNA Library Preparation kit with polyA selection. Sequencing libraries were chemically denatured in preparation for sequencing on an Illumina NovaSeq X instrument, where a 151 x 151 cycle paired end sequence run was performed using a NovaSeq X Series 10B Reagent Kit (20085594).

For assembly, first adapter trimming of de-multiplexed raw reads was performed using fqtrim (Pertea, 2015), followed by quality trimming and filtering using prinseq-lite (Schmieder & Edwards, 2011). Error correction was performed using the BBnorm ecc tool and the resulting reads were assembled using Trinity (version 2.2.1) with a k-mer length of 31 and minimum k-mer coverage of ten. Assembled transcripts were annotated using a blastx (Altschul *et al*., 1990) search (E-value setting of 1e-3) against a combined database derived from UniProt and conoserver (Kaas *et al*., 2012). Transcripts per million (TPM) counts were generated using the Trinity RSEM (Li & Dewey, 2011) plugin. Transcriptomes generated in this study were submitted to NCBI (accession number pending).

### Phylogenetic analyses

The Sanger sequenced products (and in a few cases the same genes isolated from RNAseq data) was first filtered for high-quality reads (quality threshold above 20) and aligned with a set of reference sequences from other *Conus* using MAFFT v7.526. The sequences were trimmed using trimAl v 1.2rev59 using the -gt 0.3 flag (Capella-Gutiérrez, Silla-Martínez & Gabaldón, 2009). The individual alignments were combined using AMAS (Borowiec, 2016). The combined alignmentd were used as input for IQ-TREE v2.2.2.3 with 1000 ultrafast bootstrap replicates and partitions for each gene with model selection enabled.

### Venom composition comparisons

The conotoxins were annotated as described above for each of the generated transcriptomes and a set of publicly available RNAseq dataset from conspecifics (**Table 2**). The conotoxin counts were normalized in each of the transcriptomes to a total expression of one million TPM. The counts for each superfamily were parsed and visualized in R using the package pheatmap using hieratical clustering with Ward’s D of the Euclidean distance of the natural logarithm of conotoxin gene superfamily expression.

**Table 2.** RNAseq data.

| *Species* | *Accession* | *Origin* |
| --- | --- | --- |
| *Capitaneus* | NA | Palau |
| *Distans* | NA | Palau |
| *Ebraeus* | SRR14407590 | Okinawa (Japan) |
| *Ebraeus* | SRR14407576 | Okinawa (Japan) |
| *Ebraeus* | SRR2609538 | Bali (Indonesia) |
| *Ebraeus* | NA | Palau |
| *Flavidus* | NA | Palau |
| *Leopardus* | NA | Palau |
| *Litteratus* | NA | Palau |
| *Litteratus* | SRR6378469 | Hainan (China) |
| *Litteratus* | SRR6381570 | Hainan (China) |
| *Litteratus* | SRR6381569 | Hainan (China) |
| *Magus* | NA | Palau |
| *Magus* | SRR9831255 | Ishigaki (Japan) |
| *Magus* | SRR8195628 | Philippines |
| *Miles* | SRR32023770 | Philippines |
| *Miles* | NA | Palau |
| *Milliaris* | NA | Palau |
| *Milliaris* | NA | Palau |
| *Milliaris* | SRR1544600 | Guam |
| *Milliaris* | SRR1544622 | Guam |
| *Milliaris* | SRR1544627 | Guam |
| *Milliaris* | SRR1544690 | Guam |
| *Milliaris* | SRR1544692 | Guam |
| *Milliaris* | SRR1548185 | American Samoa |
| *Milliaris* | SRR1548186 | American Samoa |
| *Milliaris* | SRR1548187 | American Samoa |
| *Milliaris* | SRR1548188 | American Samoa |
| *Milliaris* | SRR1548189 | American Samoa |
| *Milliaris* | SRR1548190 | American Samoa |
| *Milliaris* | SRR1548192 | American Samoa |
| *moreleti* | NA | Palau |
| *Musicus* | NA | Palau |
| *Mustelinus* | NA | Palau |
| *Mustelinus* | SRR32023769 | Philippines |
| *Nux* | NA | Palau |
| *Pulicarius* | NA | Palau |
| *Pulicarius* | NA | Palau |
| *Pulicarius* | SRR2609544 | Bali (Indonesia) |
| *Rattus* | SRR2609540 | Bali (Indonesia) |
| *Rattus* | NA | Palau |
| *Sponsalis* | NA | Palau |
| *Sponsalis* | NA | Palau |
| *Sponsalis* | SRR2609541 | Bali (Indonesia) |
| *Striatus* | NA | Palau |
| *Striatus* | SRR8195630 | Philippines |
| *Virgo* | NA | Palau |
| *Virgo* | SRR2608262 | Bali (Indonesia) |
| *Virgo* | SRR8195631 | Philippines |

## Results

### Sequence data and phylogeny

In total we sampled specimens belonging to 27 species. We obtained sequence data of a region of the COI gene from 34 individuals, 12S from 41, and 16S from 35 individuals.

Using maximum-likelihood methods we reconstructed the species tree of the collected *Conus* species and reference sequences. The tree shows that most specimens can be assigned to known species with few exceptions. E.g. PAL127S, has an intermediate position close to *Conus nux* Broderip, 1833, and several species show some divergence from the reference sequences, such as the clade of Palauan *Conus magus* Linnaeus, 1758.

### Venom gland transcriptomics

In addition to sequencing mitochondrial marker genes, we performed RNAseq experiments from the venom glands of 21 individuals belonging to 14 species (*Conus capitaneus* Linnaeus, 1758, *Conus ebraeus* Linnaeus, 1758, *Conus flavidus* Lamarck, 1810, *Conus leopardus* (Röding, 1798), *Conus litteratus* Linnaeus, 1758, *C. magus, Conus miles* Linnaeus, 1758, *Conus miliaris* Hwass, 1792, *Conus musicus* Hwass, 1792, *Conus mustelinus* Hwass, 1792, *C. nux, Conus pulicarius* Hwass, 1792, *Conus rattus* Hwass, 1792, *Conus sponsalis* Hwass, 1792, *Conus virgo* Linnaeus, 1758), of which *C. musicus, C. moreleti*, and *C. nux* has not previously had their venom glands sequenced.

Comparison of the venom compositions of the sequenced individuals to previously sequenced samples show a clear similarity in the overall venom composition between conspecifics based on the superfamily-level expression (**Fig. 3**). It is intriguing that *C. leopardus*, which feeds on hemichordates, has a relatively simple venom composition(Remigio & Duda, 2008) that to some degree resembles that of the fish-hunting cone snails.

**Fig 3.**
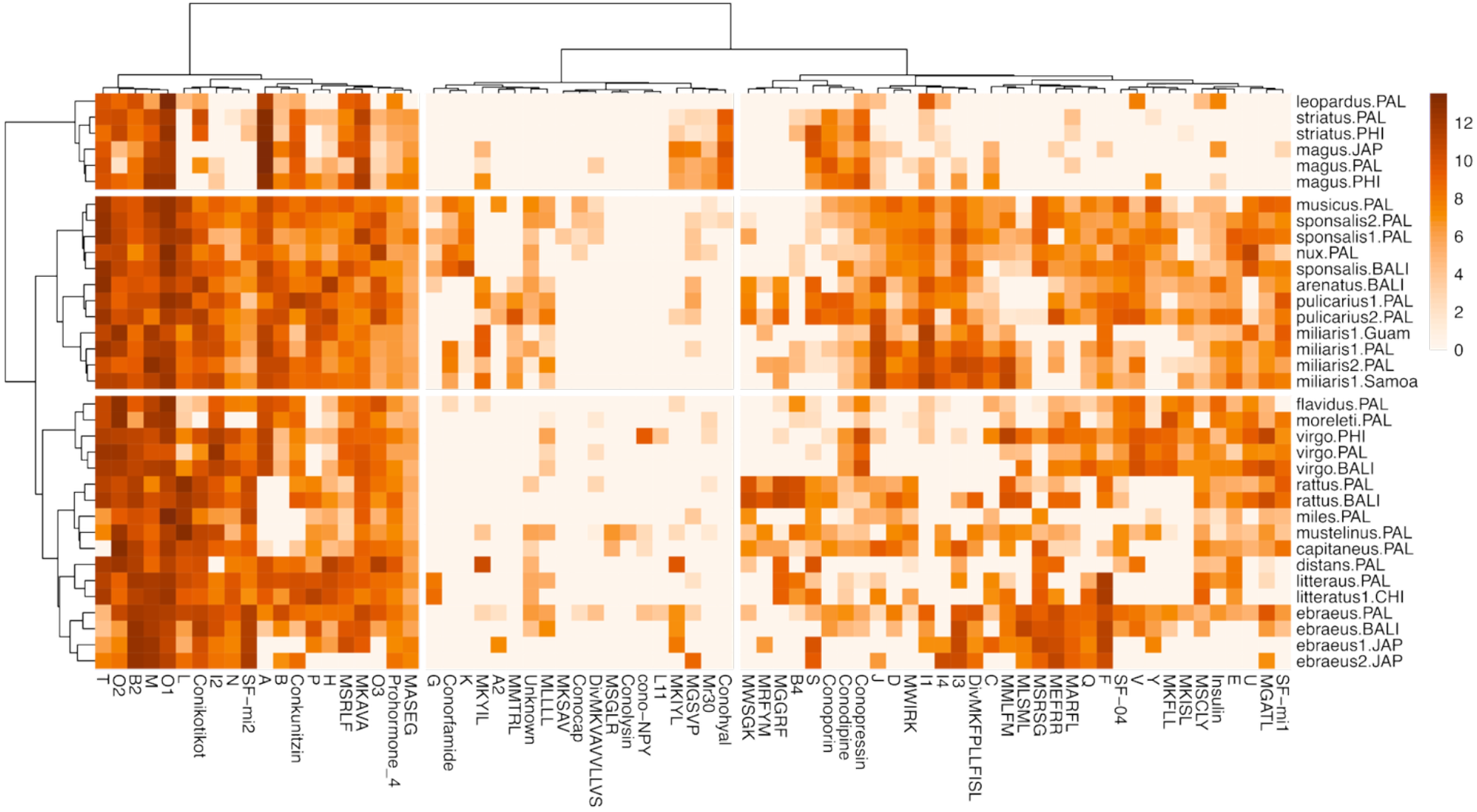
Heatmap showing venom composition in species collected in Palau, and when possible, in conspecifics. The x axis shows the different conotoxin superfamilies and the y axis shows the different transcriptomes where superfamily expression was assessed.

## Discussion

Our study presents the first integrated phylogenetic and venom-gland transcriptomic survey of *Conus* species from the archipelago of Palau, a region previously underrepresented in cone snail research. While the mitochondrial phylogeny largely confirmed the species-level identification of most sampled individuals, it also revealed some divergent lineages.

Notably, the intermediate phylogenetic position of specimen PAL127S near *C. nux* suggests the possibility of local divergence or hybridization, although the limited resolution of mitochondrial markers makes it difficult to determine its taxonomic status with confidence. Similarly, the clustering of Palauan *C. magus* sequences in a distinct clade relative to known reference sequences could indicate geographic structure or the presence of a cryptic lineage within what is considered a widespread species. Additional nuclear markers or genome-wide data would be necessary to resolve these cases.

Our transcriptomic data provide new insights into the venom composition of 14 *Conus* species in Palau, including three species, *C. musicus, C. moreleti*, and *C. nux*, for which no venom gland data were previously available. Across all sampled species, we observed consistent venom profiles at the gene superfamily level within conspecifics, supporting the hypothesis that venom composition is broadly conserved within species, even across geographic distances, although significant intraspecies variation exists between individual toxins.

Interestingly, the venom profile of *C. leopardus*, a hemichordate-feeding species, appears relatively simple compared to other species in our dataset and shows some similarity to that of fish-hunting cone snails.

These data provide a valuable baseline for future research into venom evolution, species delimitation, and the biogeography of *Conus* in the Indo-Pacific.

## Acknowledgement

We are grateful to the Republic of Palau and to Aimeliik State, Airai State, and Ngeremlengui State for permission and for the opportunity to perform the biodiversity research described in this paper. We thank the Coral Reef Research Foundation (Palau) for help with field research and logistics. The support and resources from the Center for High Performance Computing at the University of Utah are gratefully acknowledged. Sange sequencing was performed by the HSC DNA Sequencing Core at the University of Utah Health Sciences Centers using equipment provided for by the University of Utah RIF funds.

## Funding Statement

This work was supported by funding from the University of Utah.

## Conflict of Interest Statement

The authors declare they have no competing interests.

## Notes

### Competing Interest Statement

The authors have declared no competing interest.

## References

Altschul, S.F., Gish, W., Miller, W., Myers, E.W. & Lipman, D.J. 1990 Basic local alignment search tool. J Mol Biol, 215: 403–410.

Borowiec, M.L. 2016 AMAS: a fast tool for alignment manipulation and computing of summary statistics. PeerJ, 4: e1660.

Capella-Gutiérrez, S., Silla-Martínez, J.M. & GabaldÓn, T. 2009 trimAl: a tool for automated alignment trimming in large-scale phylogenetic analyses. Bioinformatics, 25: 1972–1973.

Hewitt, G.M., Johnston, A.W. & Young, J.P.W. 2013 Molecular techniques in taxonomy. Springer Science & Business Media.

Kaas, Q., Yu, R., Jin, A.H., Dutertre, S. & Craik, D.J. 2012 ConoServer: updated content, knowledge, and discovery tools in the conopeptide database. Nucleic Acids Res, 40: D325–330.

Koch, T.L., Robinson, S.D., Salcedo, P.F., Chase, K., Biggs, J., Fedosov, A.E., Yandell, M., Olivera, B.M. & Safavi-Hemami, H. 2024 Prey Shifts Drive Venom Evolution in Cone Snails. Molecular Biology and Evolution, 41.

Li, B. & Dewey, C.N. 2011 RSEM: accurate transcript quantification from RNA-Seq data with or without a reference genome. BMC Bioinformatics, 12: 323.

Olivera, B.M., Showers Corneli, P., Watkins, M. & Fedosov, A. 2014 Biodiversity of Cone Snails and Other Venomous Marine Gastropods: Evolutionary Success Through Neuropharmacology. Annual Review of Animal Biosciences, 2: 487–513.

Palumbi, S.R. 1996 PCR and molecular systematics. (No Title): 205.

Pertea, G. 2015 fqtrim: v0.9.4 release. In).

Remigio, E.A. & Duda, T.F., Jr. 2008 Evolution of ecological specialization and venom of a predatory marine gastropod. Mol Ecol, 17: 1156–1162.

Robinson, S.D. & Norton, R.S. 2014 Conotoxin gene superfamilies. Mar Drugs, 12: 6058–6101.

Schmieder, R. & Edwards, R. 2011 Quality control and preprocessing of metagenomic datasets. Bioinformatics, 27: 863–864.

